# Subsurface Microbial Colonization at Mineral-Filled Veins in 2-Billion-Year-Old Igneous Rock from the Bushveld Complex, South Africa

**DOI:** 10.1101/2024.07.08.602455

**Authors:** Yohey Suzuki, Susan J. Webb, Mariko Kouduka, Hanae Kobayashi, Julio H. Castillo, Jens Kallmeyer, Kgabo Moganedi, Amy J. Allwright, Reiner Klemd, Frederick Roelofse, Mabatho Mapiloko, Stuart J. Hill, Lewis D. Ashwal, Robert B. Trumbull

**Author notes:** Corresponding author. (Y.S).

## Abstract

Recent advances in subsurface microbiology have demonstrated the habitability of multi-million-year-old igneous rocks, despite the scarce energy supply from rock-water interactions. Given the minimal evolution coupled with exceedingly slow metabolic rates in subsurface ecosystems, spatiotemporally stable igneous rocks can sustain microbes over geological time scales. This study investigated 2-billion-year-old igneous rock in the Bushveld Complex, South Africa, where ultradeep drilling is being executed by the International Continental Scientific Drilling Program (ICDP). New procedures were successfully developed to simultaneously detect indigenous and contaminant microbial cells in a drill core sample. Precision rock sectioning coupled with infrared, fluorescence and electron microscopy imaging of the rock section with submicron resolution revealed microbial colonization in veins filled with smectite. The entry and exit of microbial cells in the veins are severely limited by tight packing with smectite, the formation of which supplies energy sources for long-term habitability. Further microbiological characterization of drilled rock cores from the Bushveld Complex will expand the understanding of microbial evolution in deep igneous rocks over 2 billion years.

## Introduction

The terrestrial subsurface is defined by depths greater than 8 m from the ground surface excluding soil [1], where a significant portion of the Earth’s prokaryotic biomass resides [2–4]. The metabolic activities of subsurface microbiomes are exceedingly slow under survival mode [5, 6], leading to an estimated turnover time ranging from several thousand to million years. Consistent with the long turnover time [7], sulfate-reducing bacteria *Candidatus* Desulforudis audaxviator endemic to the deep subsurface have undergone minimal evolution since 55–165 million years ago [8]. Similarly, minimal evolution over geological time scales has been demonstrated for deep subsurface archaeal lineages called *Candidatus* Altiarchaeota [9]. Based on the ecological and evolutionary features of subsurface microbiomes, it is hypothesized that microbes can be sustained with minimal evolution for billions of years in a geologically and tectonically stable subsurface environment.

The basement of the oceanic and continental crust is dominated by igneous rocks. Microbiological studies of the igneous basement have been intensively studied by sampling fluids from drilled boreholes [10]. In cases where collecting pristine fluid samples was technically difficult or impossible, drilled rock cores were used for microbiological characterization.

For example, in the Oman ophiolite complex, ∼100-million-year-old (Ma) mantle peridotite was drilled by the International Continental Scientific Drilling Program (ICDP). To monitor contamination, fluorescent microspheres were added to the drilling fluid and counted in the crushed and homogenized drill core samples [11]. Cells were separated from rock particles, and then DNA-stained cells with SYBR-Green I were counted by flow cytometry [12]. The cell density was up to six orders of magnitude higher at fractures and veins (∼10^7^ cells/g) than in the rock matrix (∼10^1-2^ cells/g) [11]. In another example, 100-million-year-old basaltic oceanic crust was drilled by the Integrated Ocean Drilling Program (IODP) in the South Pacific Gyre, using fluorescent microspheres in the drilling fluid for contamination control [13, 14]. In addition to contamination control using fluorescent microspheres and microbial cells extracted from the bulk rocks with ultra-low cell abundances, new visualization approaches were successfully developed to enumerate microbial cell densities at fractures and veins that exceeded 10^10^ cells/cm^3^ [13, 14].

This contribution extends the formation age of igneous rocks for microbiological investigations by targeting mafic and ultramafic rocks formed 2.05 billion years ago in the Bushveld Complex of northeastern South Africa, which is the largest mafic-ultramafic layered intrusion on Earth [15, 16]. The Bushveld drilling project of the ICDP is undertaking a 2.5-km deep drill hole in the lower zone and base of the intrusion, where ultramafic rocks are expected to possess the chemical signature of the mantle endmember of mantle-crust mixing during magma emplacement [17, 18]. Following comparatively rapid cooling of the magma chamber [19], the layered rocks have experienced minimal deformation and metamorphic alteration, providing a stable habitat for microbial life for >2 billion years. This paper reports on the first subsurface rock sample obtained from the ICDP drilling used to test new approaches for the simultaneous visualization of microbial cells and fluorescent microspheres at fractures and veins. A new spectroscopic method was also applied to obtain diagnostic spectra from single microbial cells [20]. As a result, indigenous microbial cells locally distributed along mineral-filled veins were detected.

## Materials and Methods

### Geological setting and drilling site

The Bushveld Complex is the largest mafic–ultramafic intrusion on Earth, with an areal extent of ∼500 km in the east–west direction and 350 km north–south [21]. The magmas of the Bushveld Complex were emplaced in the early Proterozoic Pretoria Group of the Transvaal Supergroup 2.05 billion years ago [15]. The entire intrusion took place in a time span within 1 million years [19]. The Bushveld ICDP drilling site is located at Marula Mine in Burgersfort (−24.50906°S, 30.08757°E), where the UG2 chromitite layer of the Upper Critical Zone is being exploited for the extraction of platinum group elements.

### Drilling and on-site core handling procedures

Rotary core barrel (RCB) drilling was conducted by Master Drilling with a diamond drill bit and drilling fluid composed of locally-sourced water mixed with a chemical additive (AMC CAP 21). Fluorescent microspheres called “Invisible Blue” (DayGlo Color Corp., pigment SPL-594NXC) with blue fluorescence under ultraviolet (UV) excitation and a particle size range from 0.25 to 0.45 μm [22] were added to the drilling fluid. The original particle concentration of ∼1×10^12^ particles/ml was diluted to a concentration of ∼1×10^9^ particles/ml.

A 30-cm long core sample with a diameter of 85 mm at a depth of 14.78 m (sample ID: 5067_3_A_010R_2_WR) was obtained on May 4^th^, 2024. After the core retrieval, the sample was laid on aluminum foil in a core box for the initial photography. The core sample was washed with drinking water filtered through a 0.2-µm-size pores (Fig. 1), and the surface was lightly flamed with a gas torch on a flame-sterilized metal tray to reduce contamination, following a procedure used in the IODP expeditions [14, 23]. After the cleaning steps, rock fractures were opened by hitting the core sample with a flame-sterilized hammer, and the rock fragments were stored at 4°C in sterilized plastic bags, then placed in vacuum-sealed plastic bags containing oxygen absorbent (Mitsubishi Chemicals Corp.).

**Figure 1.**
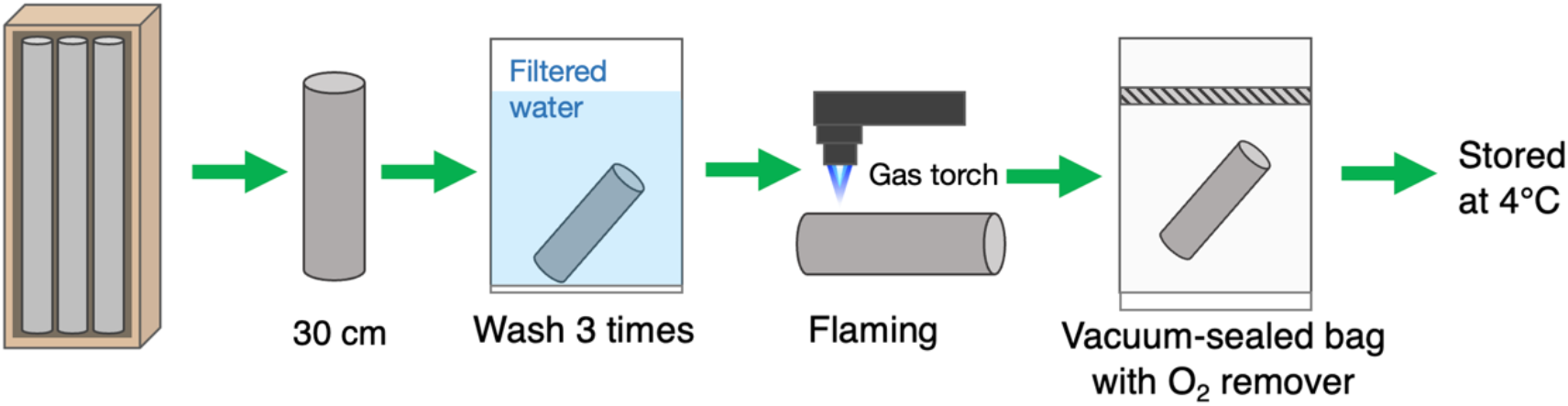
On-site handling of a drill core. A schematic illustration of the workflow for cleaning, flaming and storing of a rock core sample.

### Visualization of fluorescence microspheres in drilling fluid and the rock interior

The rock core fragments were illuminated by a 5 W UV light (Nichia Kagaku, Japan), and blue fluorescence was observed in a darkened room. After visual inspection, the rock fragment was cut using a precision diamond band saw operated without water (Meiwa Fosis Corp. DWS 3500P, Japan) in a clean booth flushed with HEPA-filtered air. A 3-mm thick section was prepared by cutting the sample perpendicular to the fracture. Fluorescence microspheres that permeated from the fracture into the rock interior were observed by a fluorescence microscope (Olympus BX51, Olympus, Japan) with a CCD camera (Olympus DP71, Olympus, Japan) and the image processing software LuminaVision (Mitani Shoji Co., Ltd.). For the drilling fluid, a fluid sample was fixed with 3.7% formaldehyde and then diluted with 10 ml filter-sterilized deionized water. After ultrasonic treatment for 30 s, a 0.2-µm-pore-size, 25-mm-diameter polycarbonate filter (Millipore) was used to collect dispersed particles. The filter was observed using the same fluorescence microscope system.

### Optical-photothermal infrared (O-PTIR) spectroscopy

The 3-mm-thick rock section was analyzed by optical-photothermal infrared (O-PTIR) spectroscopy (mIRage infrared microscope, Photothermal Spectroscopy Corp., Santa Barbara, CA, USA) using a continuous wave 532 nm laser as the probe beam. A pump beam consisting of a tunable quantum cascade laser (QCL) device (800–1800 cm^-1^; 2 cm^-1^ spectral resolution and 10 scans per spectrum) was also used. The refection mode [Cassegrain 40 objective (0.78 NA)] was used to obtain intensity maps at 1000, 1530 and 1640 cm^−1^ as well as O-PTIR spectra over the mid-IR range. Co-cultured cells of Nanoarchaeota strain MJ1 and *Metallosphaera sp*. strain MJ1HA (JCM33617) and cultured cells of *Shewanella oneidensis* (ATCC 700550) were mounted on CaF2 disks to obtain O-PTIR spectra. The Clay Science Society of Japan (JCSS) reference clay samples montmorillonite JCSS3101 ((M+.97)[Si7.8Al.02][Al3.3Fe-.2Mg.6]O20(OH)4)) and saponite JCSS3501 ((M+.98)[Si7.2Al.08][Mg6.0]O20(OH)4)) and a reference sample of nontronite coded NAu-2 ((M+.97)[Si7.57Al.01Fe.42][Al.52Fe3.32Mg.7]O20(OH)4[24] were also mounted on CaF2 disks to obtain O-PTIR spectra.

### Visualization of microbial cells in drilling fluid and the rock interior

The drilling fluid sample was treated similarly as the rock to observe fluorescent microspheres, except for the incubation of the filter in a TAE buffer containing SYBR Green I for 5 min at room temperature. This step was followed by a brief rinsing with deionized water. After O-PTIR spectroscopic analysis, the 3-mm thick section was fixed with 3.7% formaldehyde in phosphate-buffered saline (PBS) overnight and stored at 4°C in PBS. The fixed section was stained with SYBR Green I (Takara-Bio, Inc., Japan) for 30 min. After washing with ultrapure water, the section was observed using the same fluorescence microscopy system.

### Scanning Electron Microscopic (SEM) Characterization

The thin section was characterized by SEM without polishing. A field-emission-type SEM (FEI Versa 3D™) equipped with an energy-dispersive X-ray spectrometer (EDS) with a silicon drift detector (Bruker) was operated at an accelerating voltage of 20 kV.

## Results

### Evaluation of drilling fluid contamination

This study sampled a drill core sample from ∼15-m depth and a corresponding drilling fluid sample from the tank where fluorescent microspheres were added and mixed (Fig. 1B). Microscopic observations of fluorescent microspheres in the fluid sample confirmed that the concentration of fluorescent microspheres (1.1×10^9^ particles/ml) is consistent with dilution with water in the tank (Fig. 2A). In the same drilling fluid sample, DNA-stained microbial cells with a size range from ∼1 to ∼5 µm-were abundant (7.0×10^7^ cells/ml; Fig. 2B). Thus, there is a concern that permeation of the drilling fluid into the interior of rock cores could cause microbial contamination.

**Figure 2.**
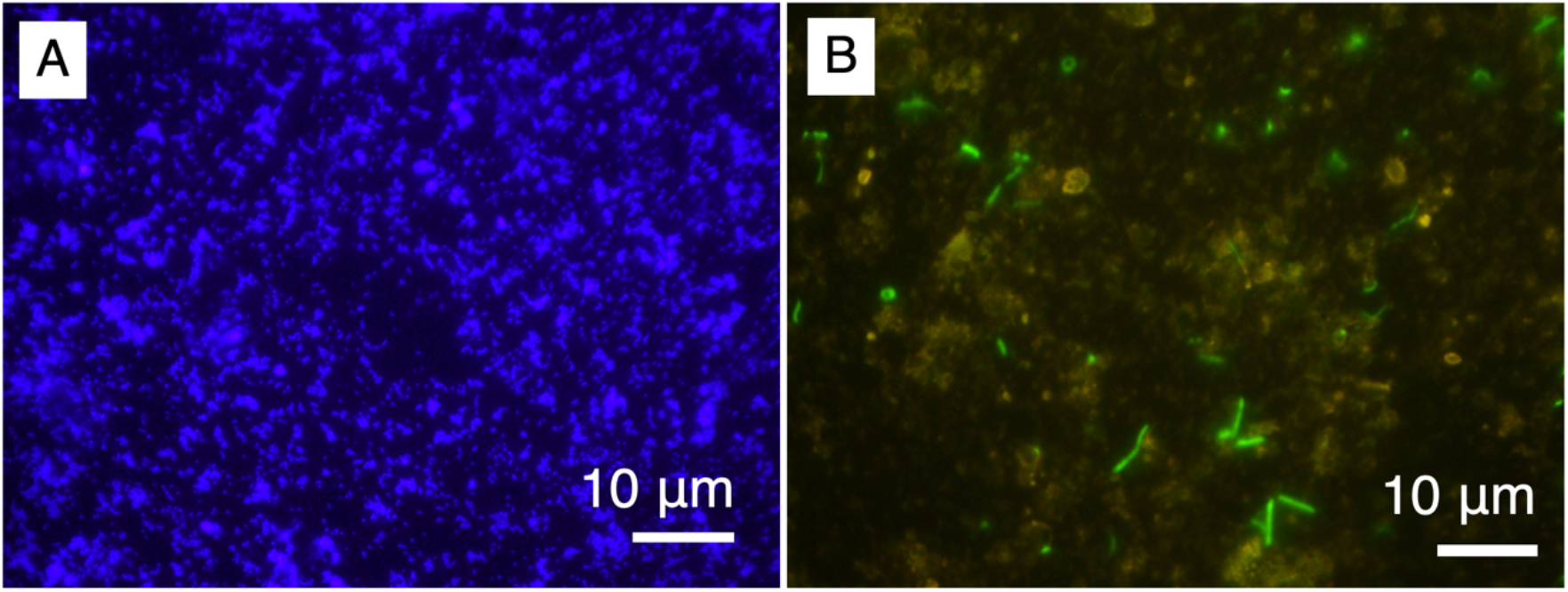
Microscopic inspection of the drill fluid sample. (A) 1000-fold magnification images of fluorescent microspheres and (B) microbial cells stained by SYBR Green I.

The drill core sample consists of norite (mafic rock with dominant plagioclase and orthopyroxene) with a porphyritic texture (Fig. 3A). After the mineral-filled fractures were opened (Fig. 3B; a yellow arrow indicates the fracture shown in Figs. 3C, E and F and Fig. 4A), a ∼10-cm long fragment was inspected for the occurrence of fluorescent microspheres by illuminating the core exterior and the fracture surface by UV light (Figs. 3C–F). Strong blue signals from fluorescent microspheres were noticed from the core exterior (Fig. 3D), whereas the blue signals were not evident at the entire fracture surface. These results demonstrate that the microspheres did not penetrate the mineral-filled fracture.

**Figure 3.**
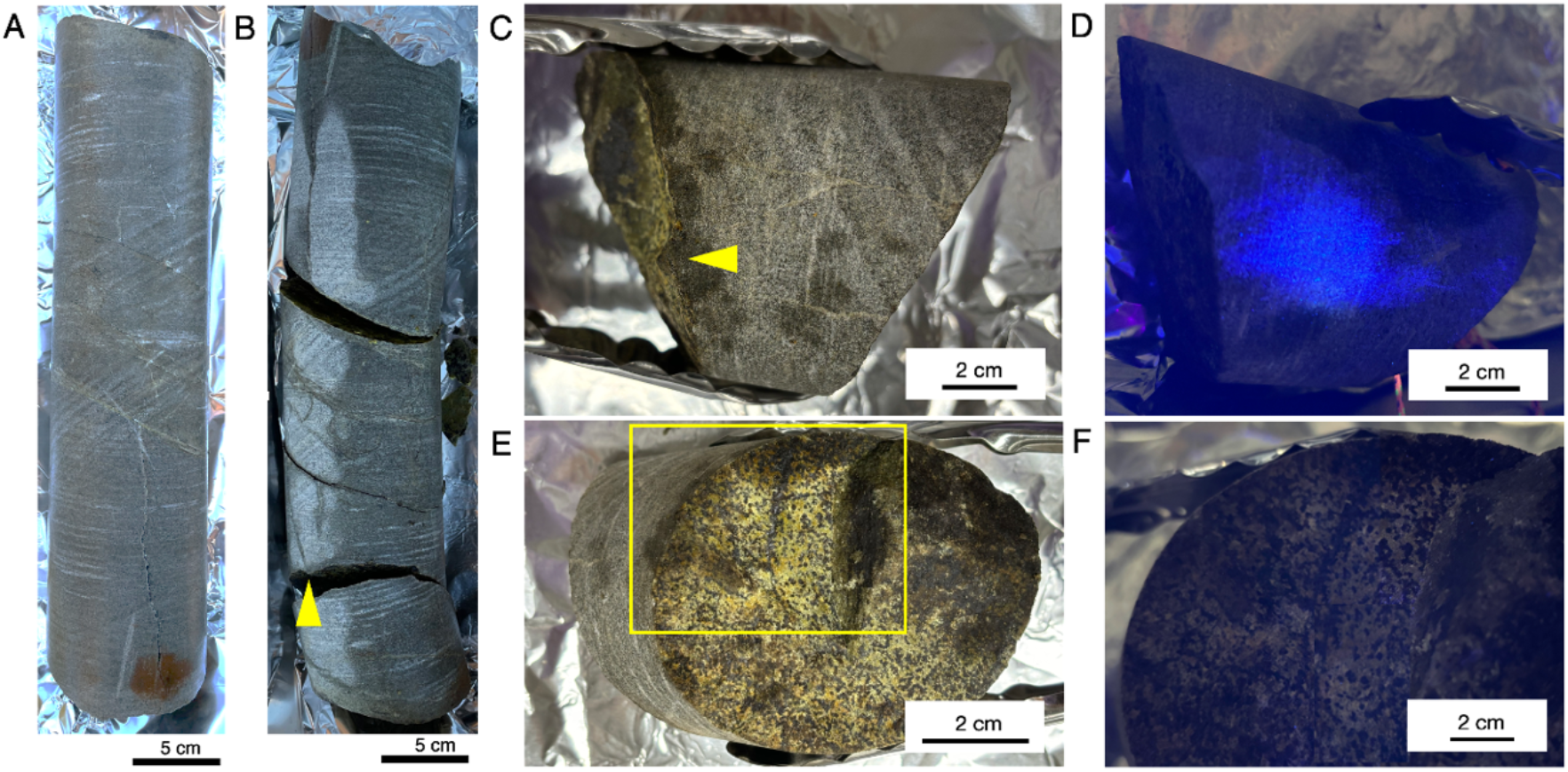
Appearance of the studied drill core sample and visual inspection of drilling fluid contamination. Photos of the cleaned whole-round core sample before (A) and after opening fractures (B). Photos of a rock fragment collected for further analysis without (C) and with (D) UV light illumination. Photos of a fracture surface without (E) and with (F) UV light illumination. Yellow arrows in (B) and (C) point to the fracture, and yellow rectangle in (E) indicates the area shown in (F).

**Figure 4.**
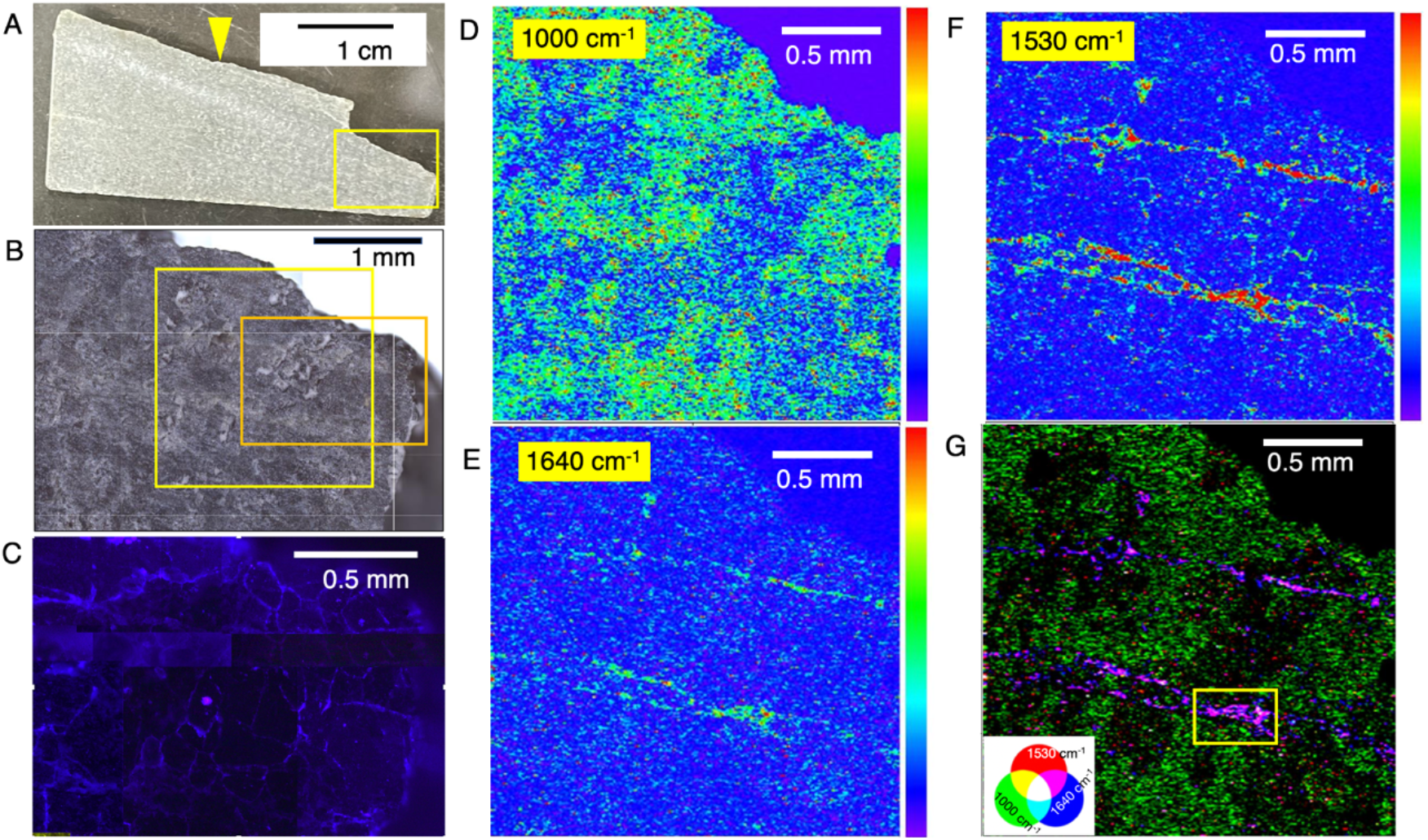
Characterization of a section from the fracture surface to the rock interior. (A) photo of the section with a yellow arrow pointing to the fracture surface. (B) microscopic image of a region of the section highlighted in a yellow rectangle in (A). (C) fluorescence microscopy image of the region highlighted by orange rectangle in (B). Intensity maps of the section at 1000 cm^-1^ (D), 1530 cm^-1^ (E) and 1640 cm^-1^ (F) obtained by optical photothermal infrared (O-PTIR) spectroscopy. The intensity maps were obtained from the area highlighted with a yellow square in (B). (G) RGB color synthesis of the three intensity maps. A yellow rectangle in (G) indicates an area where the intensity maps for RGB color synthesis was obtained in Fig. 5A.

To visualize the distribution of fluorescent microspheres from the fracture surface to the rock interior, a 3-mm thick section was examined (Fig. 4A; a yellow arrow indicates the fracture surface shown in Fig. 3E and F). Microscopic observations revealed that fluorescent microspheres were not detected from the surface to the interior of the section (Fig. 4B and C). These results also demonstrated that in addition to the rock surface, the rock interior was not contaminated by the drilling fluid. This new approach for contamination evaluation was successfully developed for the BVDP project.

### Microbial cell detection from the rock interior

Before staining in SYBR-Green I solution, which causes organic contamination, the detection of microbial cells was performed by a nondestructive technique called O-PTIR spectroscopy (Fig. 4D–G). Three wave numbers were selected for intensity mapping, whereby 1000 cm^-1^ and the combination of 1530 and 1640 cm^-1^ indicate the presence of silicate minerals and microbial cells, respectively [25, 26]. Mapping of a 2×2 mm area revealed veins nearly parallel to the fracture with high intensities at 1530 and 1640 cm^-1^ (Fig. 4E and F), whereas the ubiquity of silicate minerals was shown by the 1000 cm^-1^ map (Fig. 4D). From one of the vein-like regions (400×250 μm shown as a yellow rectangle in Fig. 4G), RGB mapping was performed to resolve the intensity distributions of 1000, 1530 and 1640 cm^-1^ (Fig 5A). Although the 1640 cm^-1^ intensity was strong throughout the vein region, the high intensity at 1530 cm^-1^ appeared spotty and partly overlapped with the high intensity of 1000 cm^-1^ (shown in white color in Fig. 5A). Mapping of white spots at the finer resolution showed a micrometer-scale heterogeneity (Fig. 5B). In Fig. 4C, O-PTIR spectra obtained from the vein-like region represented by Point 1 (Fig. 5B) had two major peaks attributed to SiO2 (∼1000 cm^-1^) and H2O (∼1640 cm^-1^). The overall spectra are similar to that of saponite, a smectite mineral commonly formed by low-temperature alteration of mafic and ultramafic rocks [13, 27]. Several O-PTIR spectra from the white spots contained two peaks of ∼1530 and ∼1640 cm^-1^ attributed to amides I and II, which are diagnostic for proteins in microbial cells (Points 5 and 6 in Fig. 5C). The rest of the peaks in the O-PTIR spectra are similar to those obtained from adjacent points (Points 3 and 4) and a distant point (Point 2), suggesting the coexistence of some mineral phases with microbial cells.

**Figure 5.**
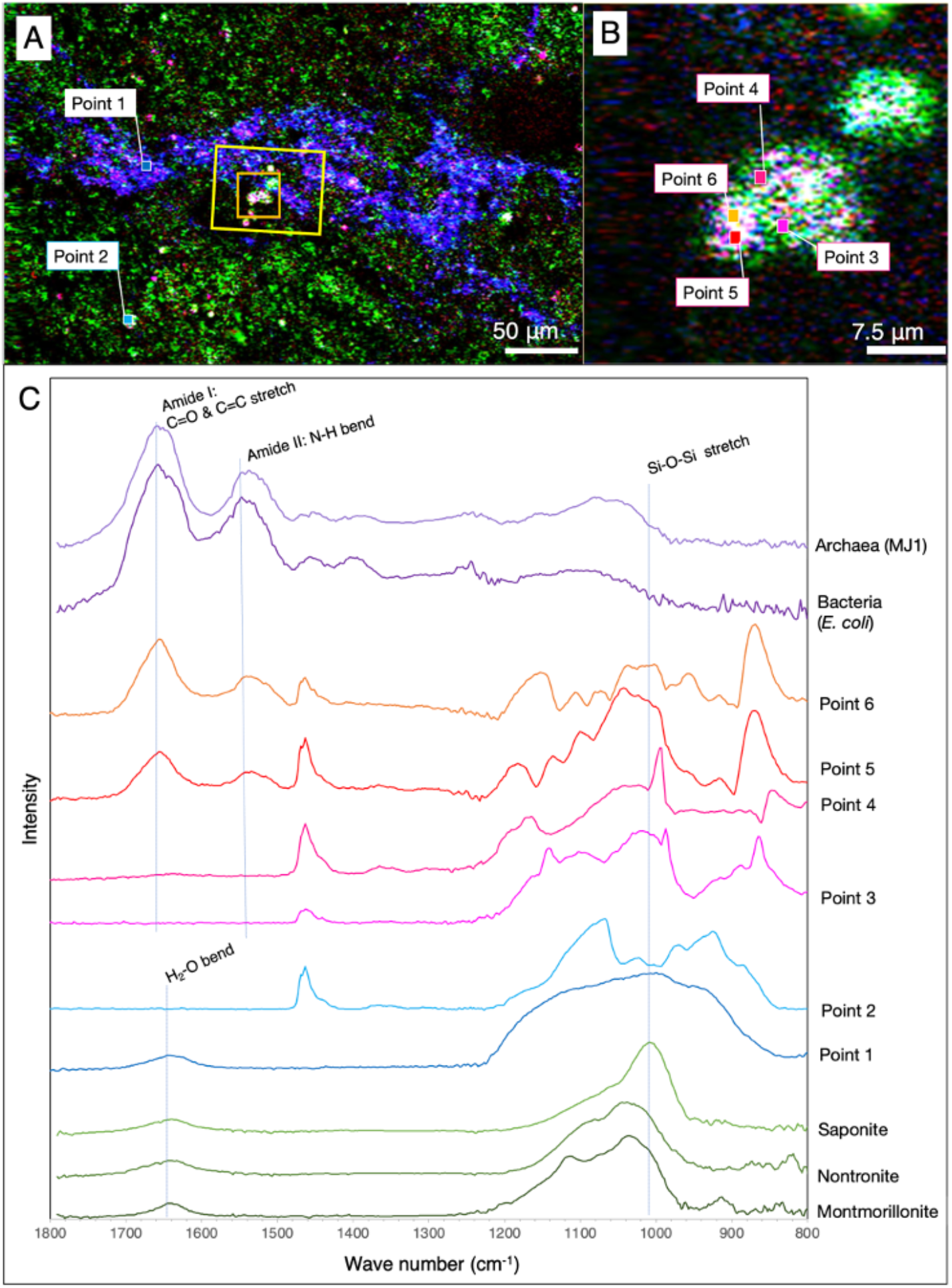
Characterization of a vein-like region in a section. RGB color synthesis of the three intensity maps at 1000, 1530 and 1640 cm^-1^ from areas highlighted by a yellow rectangle in Fig. 4G (A) and highlighted by an orange square in Fig. 5A (B). O-PTIR spectra from Points 1 and 2 in Fig. 5A and from Points 3-6 in Fig. 5B, cultured cells of *Nanobdella aerobiophila* strain MJ1^T^ (= JCM33616^T^) and *Metallosphaera sedula* strain MJ1HA for an archaeal reference and *Shewanella oneidensis* strain MR-1^T^ for a bacterial reference and smectite references of saponite, nontronite and montmorillonite (C).

After obtaining the O-PTIR spectra, the section was stained in the SYBR-Green I solution. Fluorescent microscopic observations showed that greenish signals were obtained from the veins indicated by pink arrows in Fig. 6A. High-magnification observations of the white spots revealed that the greenish signals were morphologically similar to microbial cells with a size range less than 1 μm (Fig. 6B). Taking the amide peaks in the O-PTIR spectra into account, we conclude that the veins are colonized by microbes. In addition, the morphological difference in microbial cells from the vein and the drilling fluid (Fig. 2B) indicates that the microbial cells found in the vein are indigenous rather than originating from contamination.

**Figure 6.**
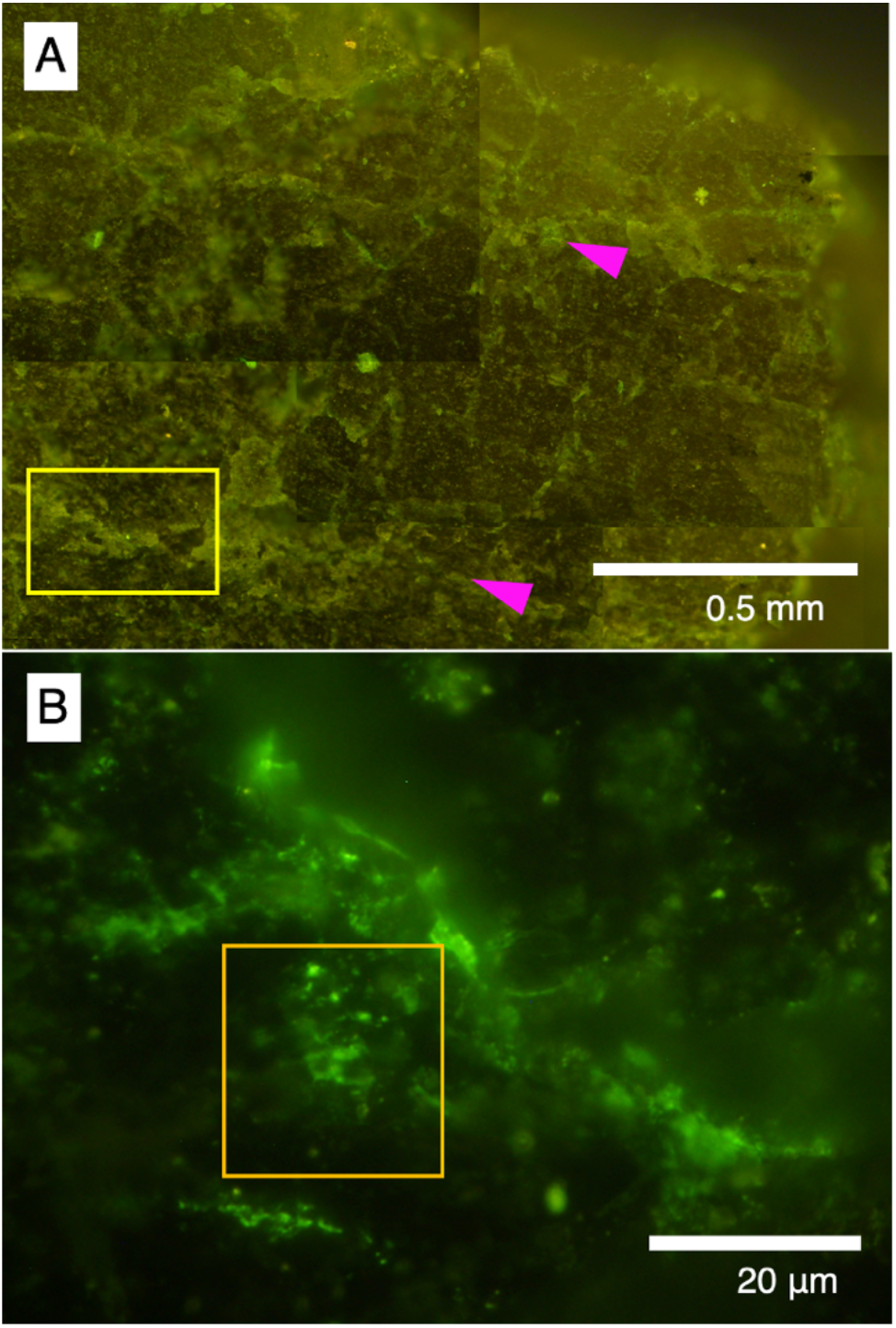
Visualization of microbial cells after SYBR-Green I staining. (A) Fluorescence microscopic images of the large area from the section highlighted by an orange rectangle in Fig. 4B, and (B) the small area highlighted by a yellow rectangle in Fig. 5A. The yellow rectangle in (A) is the same as shown in Fig. 5A, and the orange square in Fig. 6B is the same as in Fig. 5B. Pink arrows in (A) point to the veins shown in Fig. 4.

### Mineralogical characterization of the vein with microbial colonization

After the DNA-stained microbial cells were visualized by fluorescence microscopy, the section was analyzed by SEM-EDS to determine the vein-fill mineralogy (Fig. 7). From a region associated with microbial cells in the vein (Fig. 7A), Si, O, and Mg were detected as major elements (Fig. 7B). A ratio of peak intensities of Si and Mg was ∼2:1, suggesting that the vein is filled with a 2:1 tri-octahedral clay mineral. The presence of water molecules in the O-PTIR spectrum (Point 1 in Fig. 5C) and the low intensity of interlayer Ca in the EDS spectrum (Fig. 7B) are consistent with the occurrence of saponite [13].

**Figure 7.**
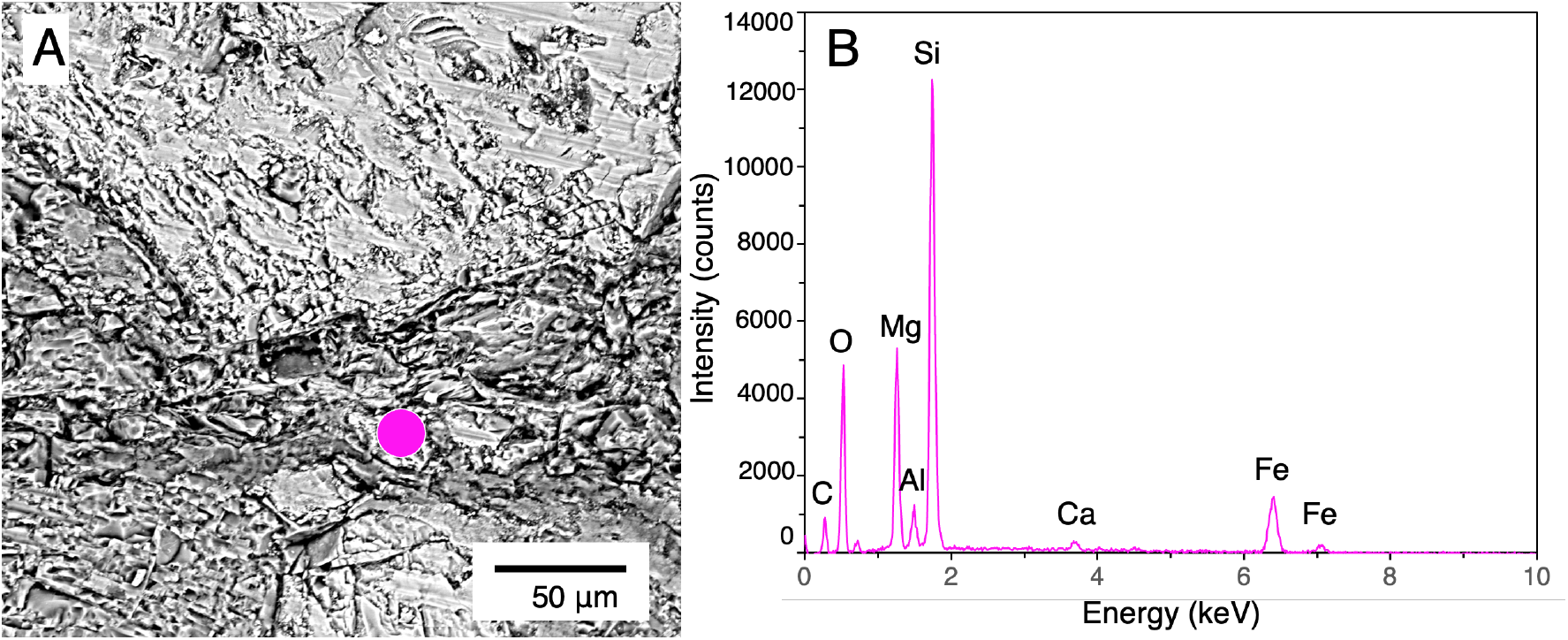
Mineralogical characteristics of the vein associated with microbial colonization. (A) A back-scattered electron image of the area in the section, where microbial colonization is shown in Fig. 6B. (B) an EDS spectrum from the pink circle in (A).

## Discussion

### Technical advances in microbial detection from drilled igneous rocks

For microbiological investigations of igneous rocks in the subsurface, the use of fluorescent microspheres started in terrestrial drilling [28] and was later adopted to ocean drilling [29]. A common procedure for detecting fluorescent microspheres is to crush the rock core material into powder. Fluorescent microspheres are extracted from the powdered material by suspending the sample in 3% NaCl solution and collecting fluorescent microspheres from the supernatant. This procedure cannot distinguish the exact location of fluorescent microspheres or the proportion of microspheres in fractures/veins and in the rock matrix. The direct observation of fluorescent microspheres developed in this study enables the location-specific detection of fluorescent microspheres in fractures/veins.

The new microbial detection approach also differs from previous approaches, in which rock fragments were embedded in hydrophilic resin to prepare thin sections [13, 14]. Although the resin can preserve rock features found in the void space [30], fluorescent microspheres are removed during dehydration of the fragments with ethanol and subsequent resin embedding. Dry cutting with a diamond band saw has technical advantages in avoiding analytical interferences for organic compounds.

In the previous approaches, staining of microbial cells with SYBR Green I was performed using ∼100-μm thick section [13, 14, 30]. To verify the greenish signals of SYBR Green I stained microbial cells or materials strongly adsorbing SYBR Green I [31], 3-μm thick sections were prepared by focused ion beam (FIB). Subsequent nanoscale secondary ion mass spectrometry (NanoSIMS) measurements were performed to show the co-localization of ions, such as ^32^S^-, 31^P^-^ and ^12^C^14^N^-^ [14, 30]. The FIB section fabrication for NanoSIMS analysis was limited to very small areas (∼10×10 μm), whereas mapping at the millimeter scale down to the tens of micrometer scale can be performed by O-PTIR spectroscopy.

### Habitability of the 2-billion-year-old igneous rock

In∼100-Ma igneous rocks obtained from drill cores from the ICDP Oman ophiolite drilling and from IODP Expedition 329 to the South Pacific Gyre, microbial colonization was found to be associated with fractures and veins [11, 13, 14]. In the IODP basaltic basement study, the association of microbial cells and smectite was clearly demonstrated [13, 14]. Smectite can effectively adsorb organic matter [32] and organic matter is also produced on smectite *in situ* [33]. Furthermore, it is known that H2 is derived from H2O reacted with Fe(II) in the transformation of silicate minerals into smectite [34]. In the case of the 2-billion-year-old igneous rocks investigated in this study, veins with microbial colonization are associated with smectite. The tight sealing of the veins with smectite prevented the contamination of microbial cells from the drilling fluid. In turn, indigenous microbes are immobile and survive in the veins by metabolizing inorganic and/or organic energy available around smectite. The age of vein formation and smectite mineralization needs to be determined for the duration of habitability.

## Conclusion

Combining previously developed procedures using fluorescent microspheres and SYBR Green I for contamination control and microbial detection, respectively, this study successfully developed new procedures for the detection of microbial colonization in fractures/veins in drill cores of igneous rocks, involving precision sectioning by a diamond band saw and single-cell level detection by IR spectroscopy. Application of the new procedures to 2-billion-year-old rocks from the Bushveld Complex revealed that indigenous microbes are colonizing veins filled with smectite. Future studies are directed to clarify the phylogenomic and metabolic profiles of the microbiome and the formation history of fractures/veins in the rock interior.

## Acknowledgments

We are grateful to Master Drilling for the drilling operation. The authors would like to acknowledge Mpho Molautsi, Katja Heeschen, and Kwena Mathopa for on-site assistance. YS and JHC are funded by the Japan Society for the Promotion of Science (JSPS)/National Research Foundation (NRF) Bilateral Joint Research Project. The Bushveld Drilling Project (BVDP) receives funding from the International Scientific Drilling Program, the National Research Foundation of South Africa, the Deutsche Forschungsgemeinschaft (DFG) of Germany and the South African Council for Geoscience. The authors would like to thank Enago (www.enago.jp) for the English language review.

## Author contributions

Frederick Roelofse, Reiner Klemd, Lewis Ashwal, Susan Webb and Robert Trumbull contributed to the successful application and implementation of the Bushveld ICDP project. Julio Castillo, Jens Kallmeyer, Kgabo Moganedi, Stuart Hill, and Amy Allwright were involved in preparing and performing on-site sample collection. Frederick Roelofse and Mabatho Mapiloko described the geology of the rock sample examined. Mariko Kouduka, Hanae Kobayashi, and Yohey Suzuki conducted experimental work, data collection, and analysis. The first draft of the manuscript was written by Yohey Suzuki who played a coordinating role in the study and provided the final approval of the manuscript. All authors contributed to writing and editing of the manuscript.

## Data availability

The datasets analyzed during the current study are available from the corresponding, first and second authors upon reasonable request.

## Ethics approval

This is an observational study and does not involve human or animal subjects, therefore ethics approval was not required.

## Consent to participate

This is an observational study and does not involve human or animal subjects, therefore no consent to participate was necessary.

## Consent to publish

This is an observational study and does not involve human subjects, therefore no consent to publish was required.

